# Anthocyanin biosynthesis gene activation in nitrogen deprived *Utricularia gibba* L. under light or darkness

**DOI:** 10.64898/2026.08.27.747637

**Authors:** Samuel Nestor Meckoni, Julie Anne V.S. de Oliveira, Boas Pucker

## Abstract

*Utricularia gibba* L. is an aquatic carnivorous plant with a diverse set of capabilities. Reddening of traps frequently occurs in old *in vitro* cultures. While anthocyanins are often responsible for red coloration in plants, not every plant turns red. Stress factors like high light or excess sucrose have previously been shown to induce the formation of anthocyanins. Here, we hypothesized the red trap formation to be dependent on nutrient deprivation and tested nitrogen deprivation. The results suggest, that only in combination with light, nitrogen deficiency leads to the activation of the complete anthocyanin biosynthesis pathway and visible red coloration. However, in darkness, anthocyanin biosynthesis appears generally less active compared to light conditions and expression of most anthocyanin biosynthesis genes is not significantly upregulated under nitrogen deficiency.

**Graphical Abstract:** 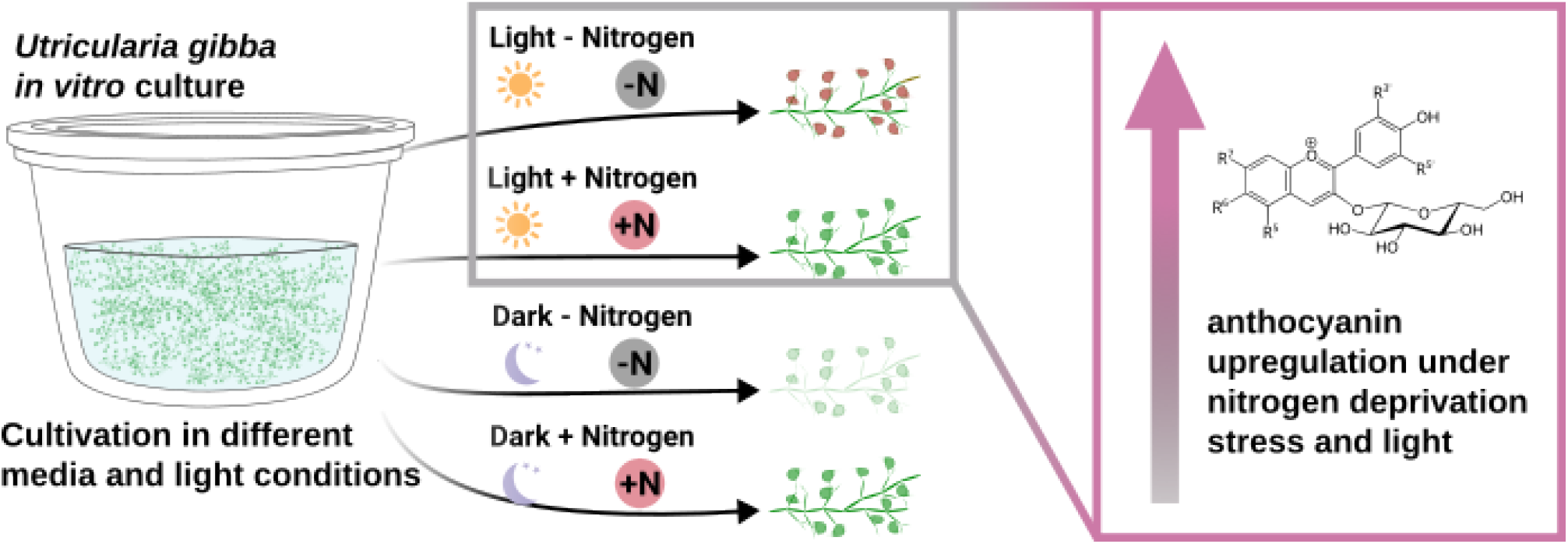

## Introduction

Plants are sessile and need to cope with diverse abiotic and biotic stresses such as high light intensities, nutrient starvation, and pathogenic microorganisms. One resilience mechanism of plants is the production of specialized metabolites, also referred to as secondary metabolites, as their functions are usually believed to not always be crucial to plants survival under physiological conditions (Weng 2014). A diverse group of specialized plant metabolites are the flavonoids, with roles such as coloration, or UV protection, especially for the subclass anthocyanins (Winkel-Shirley 2001; Grünig et al. 2025). A specialized role of anthocyanins is their accumulation and increased fitness under nitrogen deficiency stress, which has been shown to be connected to high light conditions (Shi and Xie 2010; Liang and He 2018; Wolff et al. 2026). Derived from phenylalanine through the phenylpropanoid pathway, anthocyanins are formed by the activity of multiple enzymes of which the first ones are shared with other flavonoids (Winkel-Shirley 2001). Anthocyanin-specific enzymes include: dihydroflavonol 4-reductase (DFR), anthocyanidin synthase (ANS), anthocyanin-related glutathione S-transferase (arGST), UDP-dependent anthocyanidin 3-O-glucosyltransferase (UGT), with more diverging specialized decorating enzymes downstream (Grünig et al. 2025; Choudhary et al. 2026). Regulation of the anthocyanin biosynthesis is controlled by the MBW complex comprizing a MYB, a bHLH, and a WD40 transcription factor protein (Ramsay and Glover 2005; Gonzalez et al. 2008). In particular, subgroup 6 members of the R2R3-MYB gene family have been attributed to activate anthocyanin biosynthesis (Stracke et al. 2001; Borevitz et al. 2000; Grünig et al. 2025). Additionally, subgroup 4 R2R3-MYBs have been attributed to a repressor role in connection with the anthocyanin biosynthesis (LaFountain and Yuan 2021).

*Utricularia gibba* L. is an aquatic carnivorous plant, belonging to the family Lentibulariaceae (Taylor 1994). Remarkably, all genera in this plant family are carnivorous: *Pinguicula* has sticky leaf traps, *Genlisea* fish trap like structures, and *Utricularia* negative pressure driven suction bladder traps (Müller et al. 2004). While sticky leaf traps have been reported in many different plants, representing a striking example of convergent evolution, the *Genlisea* and *Utricularia* specific trap mechanisms have only been found in these genera respectively (Ellison and Gotelli 2009; Müller et al. 2004). Especially the suction bladders of *Utricularia* have fascinated scientists for more than a century (Darwin 1875). They consist of a thin cell layer surrounding an extracellular lumen, which is enclosed from the external environment by a trap door sealed with mucilage and cuticle wax (Lloyd 1942; Vincent et al. 2011). When triggered, the trap door buckles inward of the trap for a short moment, before closing the bladder again, letting surrounding water and putative prey inside (Vincent et al. 2011). To regain their negative pressure, water and ions are pumped out, while retaining prey inside (Sydenham and Findlay 1975). The trap biology includes additional features, like spontaneous trap action without an external trigger, diverse microbial communities living inside the traps, as well as the supply of photosynthetically-fixed carbon to the traps (Płachno et al. 2012; Sirová et al. 2010). Collectively, *Utricularia* are not only an excellent model system to study their unique traps, but specifically these make it an ideal candidate for diverse research questions regarding evolution, (micro)ecology as well as the application of nature-derived inspirations for synthetic biology and biotechnology (Sirová et al. 2018). Specifically *U. gibba* has been used for genetic engineering before and due to its small size and fast growth capabilities, it can be considered an *Utricularia* model plant (Bushell 2016; Lee et al. 2019; Oropeza-Aburto et al. 2020; Płachno et al. 2026; Whitewoods 2020; Meckoni 2024).

Here, we report a genome sequence of *U. gibba* derived from an *in vitro* culture as well as its behavior during nitrogen deficiency stress under variant light conditions regarding anthocyanin biosynthesis gene regulation.

## Results

The newly generated genome sequence of *U. gibba* has a total length of 118,553,830 bp with a contig N50 of 5,593,449 bp. The BUSCO score is 91.0% and 85.6% (with lamiales_odb12 and lamiales_odb12.2, respectively) and the LAI score 12.22. In total, 24,590 protein coding gene models have been structurally annotated, with a BUSCO score of 90.6% and 85.3% (with lamiales_odb12 and lamiales_odb12.2, respectively). Previously, it was observed that the traps of *U. gibba in vitro* cultures become red after several weeks in the same cultivation media (Figure 1, Meckoni (2024)).

**Figure 1.**
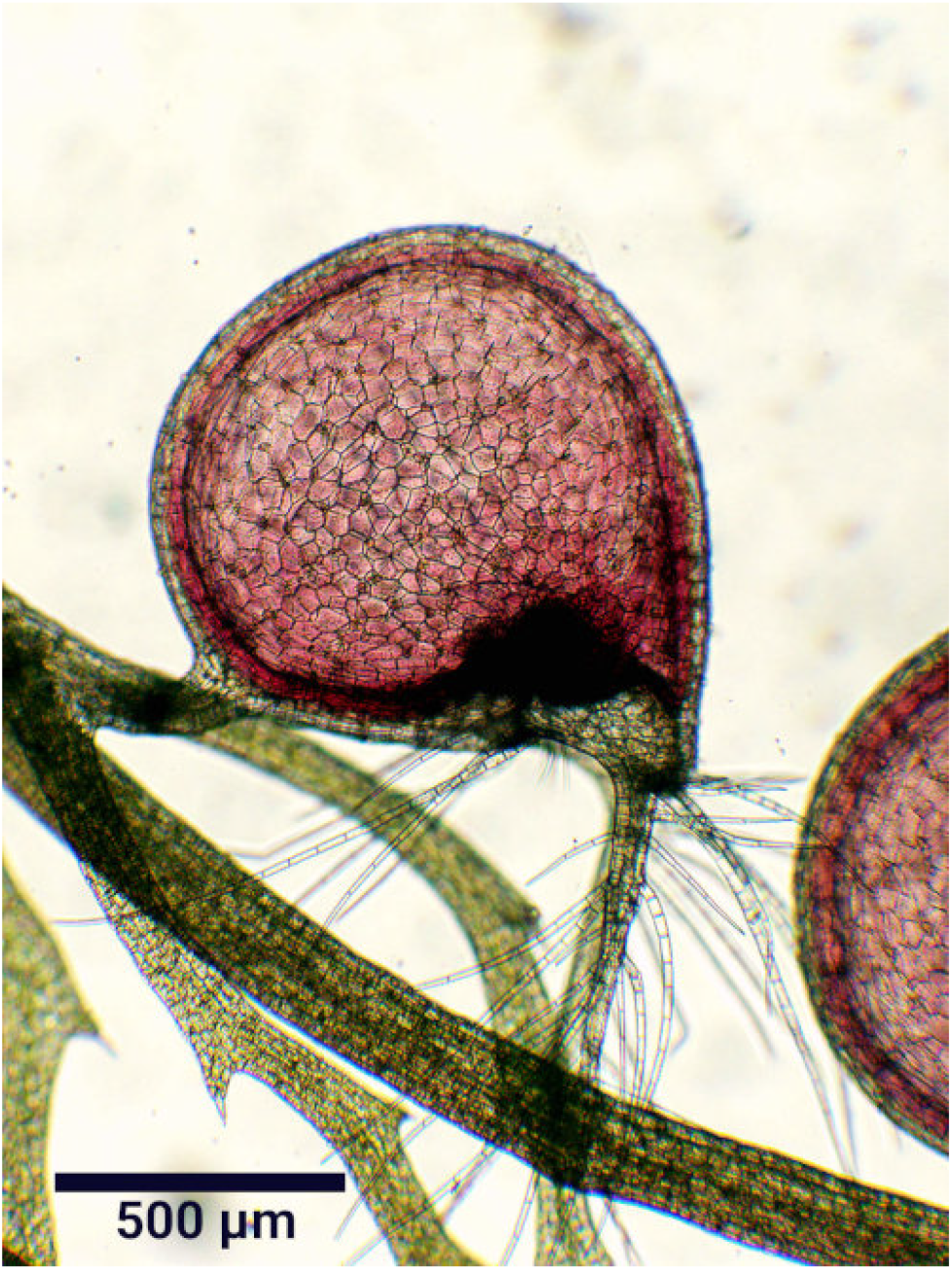
Microscopic bright field image of a trap of an Utricularia gibba in vitro culture that has been in the same media for at least ca. two months. The scale bar represents 500 µm.

The nitrogen deprivation experiments resulted in particularly green tissue for nitrogen containing samples (Additional file 1 and Additional file 2). For the nitrogen deficiency samples, only a few plant parts appear green, and many whitish (Additional file 3 and Additional file 4). Remarkably, only the samples under nitrogen deprivation and light conditions show clear reddening of many parts of the tissue, most pronounced in the traps, but also visible in other tissue parts (Additional file 3).

Relevant gene models for the biosynthesis of flavones, flavonols and anthocyanins have been annotated (Additional file 5, Additional file 6). Starting from the phenylpropanoid pathway, there are three candidates of *PAL*, one of *C4H*, and two of *4CL*, that have been found with a 100% conserved residue score and a high similarity score (>90%). For the following genes, one best candidate was identified: *CHS, CHI, F3H, F3’H, FLS, DFR, ANS, arGST*, and *UGT*; as well as two for *FNSII*. For all these genes, the conserved residue score was 100%, except for *CHS* (97%), *ANS* (96%), and *UGT* (97%). Across the stress experiment samples, the anthocyanin biosynthesis specific genes are all differentially expressed (Figure 2). In light/dark conditions, *CHI, F3H, F3’H, DFR*, and *arGST* are significantly more expressed under nitrogen deprivation conditions, while this is only true in light conditions for *CHS, ANS*, and *UGT*. By contrast, arGST is more highly expressed in light compared to dark conditions. Albeit some genes from *F3H* onward are similarly differentially expressed between the light/dark conditions, the gene expression appears generally lower in dark conditions (Figure 2). Non-anthocyanin specific genes show a different pattern. One *FNSII* candidate, daUtrGibb.v01.v01.ptg000005l.g013910.1, shows diametrical expression changes between normal to nitrogen deprivation, being significantly more expressed in nitrogen deprivation under light, while being less highly expressed in nitrogen deprivation under darkness. The other *FNSII* candidate, daUtrGibb.v01.v01.ptg000005l.g013960.3, shows upregulation under nitrogen deprivation under both, light and dark conditions. In total, 123 gene models were annotated as members of the MYB transcription factor family (Additional file 7). Out of these, candidates known to be related to the up- or downregulation of the flavonoid and anthocyanin biosynthesis have been compiled in a heatmap (Additional file 8). They comprise diverse expression patterns, ranging from virtually no expression (e.g.: daUtrGibb.v01.v01.ptg000002l.g006200.1) to highly different differential expression (e.g.: daUtrGibb.v01.v01.ptg000036l.g012070.1, Additional file 8).

**Figure 2.**
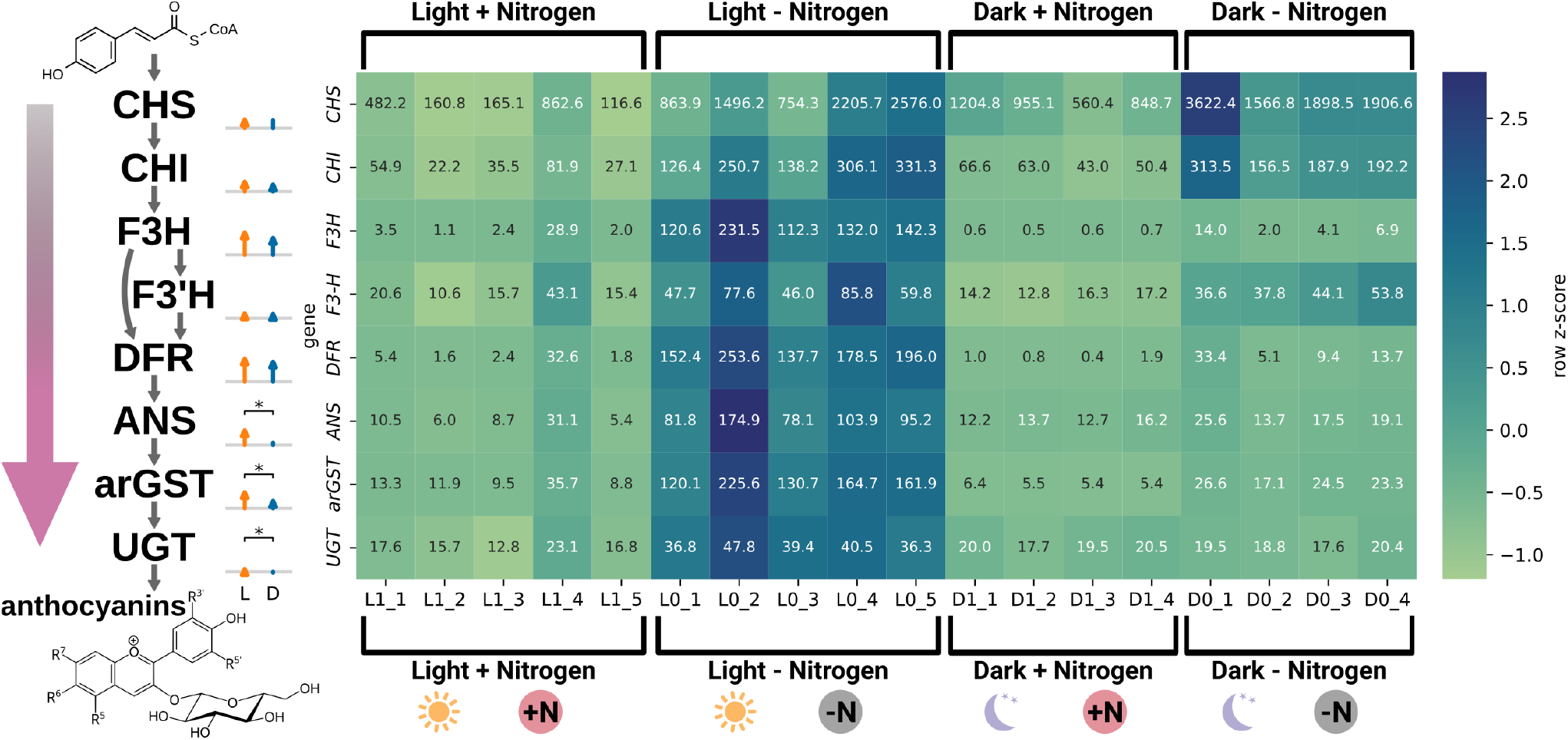
Expression pattern of structural anthocyanin biosynthesis genes under nitrogen starvation in the presence or absence of light. The heatmap is based on kallisto TPMs, and the color is z-score normalized by gene/row. The bar charts next to the heatmap show the log2 fold change between nitrogen containing (+N) and nitrogen starvation (-N) media for normal light conditions (orange bars, L) and dark conditions (blue bars, D). If the change in gene expression between the different media conditions is significant (padj ≤ 0.05), the bar is enriched by an arrow. The bracket with an asterisk marks a significant difference (padj ≤ 0.05) in the log2 fold change between the different light conditions (DESeq2 interaction). Gene/protein abbreviations: CHS = chalcone synthase, CHI = chalcone isomerase, F3H = flavanone 3-hydroxylase, F3’H = flavonoid 3’-hydroxylase, DFR = dihydroflavonol 4-reductase, ANS = anthocyanidin synthase, arGST = anthocyanin-related glutathione S-transferase, UGT = UDP-dependent anthocyanidin-3-O-glucosyltransferase. The illustrative chemical formulas represent the common precursor coumaroyl-coenzyme A (top structure) and a generalized form of anthocyanidin 3-O-glucoside (bottom structure). The corresponding daUtrGibb.v01.v01 IDs can be obtained from Additional file 9.

## Discussion

This study elucidates the influence of nitrogen deprivation and light on the biosynthesis of anthocyanins in *U. gibba*. Our findings suggest that light is required to trigger the formation of anthocyanins. After the initial observation of *U. gibba* traps turning red when grown for a long time in the same cultivation media (Figure 1), it was suggested for two reasons that this might have been connected to nutrient deprivation. First, after a longer time period the plant has used most of the available nutrients, until the nutrient level resembles a stress situation for the plant. Second, during this time period, the plant grew, and thus had became more resource-consuming. Since nitrogen availability is very important for plants and often the bottleneck of efficient growth, the hypothesis arose that nitrogen deficiency causes the formation of the red traps (Vitousek and Howarth 1991; Wolff et al. 2026). To further test the influence of light, the present experiment was conducted. After establishing an improved new contiguous genome sequence of *U. gibba* and a corresponding functional annotation of gene models, all relevant genes for the formation of anthocyanins could be identified in *U. gibba* (Figure 2, Additional file 8). The transcriptomics data revealed, that the difference in expression between control and the nitrogen deficiency is not significantly different between light and dark conditions for the flavonoid biosynthesis genes *CHS, CHI, F3H*, and *F3’H* (Figure 2). Albeit the overall expression appears lower for *F3H*, the nitrogen stress seems to have the same effect on the expression of these genes. This is in strong contrast to the downstream anthocyanin specific biosynthesis genes. Except for *DFR*, the change in expression between the control and the nitrogen deficiency is significantly higher under light conditions, compared to dark conditions. The overall expression for *DFR* is higher under light compared to dark conditions. This concludes that the overall flavonoid biosynthesis is upregulated under nitrogen deprivation, and within that, specifically the anthocyanin biosynthesis when light is present. In competition with the anthocyanin formation through F3H, flavones can be produced through FNSII. Two *FNSII* candidates found in *U. gibba* showed an expression pattern that suggests flavones are formed under both, light and dark conditions. Since light is required for nitrogen deprivation to stimulate anthocyanin biosynthesis, the resulting increase in anthocyanin flux may occur at the expense of flavones. Thus, flavones might be comparatively increased in the dark nitrogen deprivation conditions.

For the activation of the anthocyanin biosynthesis genes, MYB transcription factors are crucial (Marin-Recinos and Pucker 2024). Subgroup 6 members of the R2R3-MYB gene family are specifically known to activate the anthocyanin biosynthesis (Stracke et al. 2001; Borevitz et al. 2000; Grünig et al. 2025). Multiple subgroup 6 members have been found in *U. gibba*, all of them being categorized as orthologs of a known subgroup 6 MYB gene of *Arabidopsis thaliana* (Additional file 8). Of these, specifically daUtrGibb.v01.v01.ptg000002l.g006150.1 shows a high expression under light, suggesting that it is light-dependent. Another ortholog, daUtrGibb.v01.v01.ptg000016l.g004590.1 is highly expressed in light and also more highly expressed in nitrogen deprivation conditions (Additional file 8). These MYB transcription factors are the best candidates for the strong activation of the anthocyanin biosynthesis under nitrogen deprivation in light. In contrast, members of the subgroup 4 of the R2R3-MYB gene family are connected to anthocyanin biosynthesis repression. Two subgroup 4 MYBs found in *U. gibba* have an expression pattern that can explain the absence of upregulation of the anthocyanin biosynthesis under nitrogen deprivation in dark conditions: daUtrGibb.v01.v01.ptg000039l.g005440.1 and daUtrGibb.v01.v01.ptg000005l.g000340.1, both classified as orthologs of the *Arabidopsis thaliana* gene At2g16720-At2R-MYB007 (Additional file 8). However, repressors alone might not be responsible for the low expression levels of anthocyanin biosynthesis genes as subgroup 6 MYBs are less highly expressed as well. Both, upregulated repressors and downregulated activators could explain a lack of anthocyanin biosynthesis under dark conditions. A lack of nitrogen might lead to less efficient growth, while the photosynthesis machinery continues to work normally. Thus, the excess of fixed carbon needs to be channeled away and due to their sugar moieties, anthocyanins have been suggested to act as a carbon sink (Grünig et al. 2025; Jezek et al. 2023; Lo Piccolo et al. 2018). In contrast, anthocyanins may act as a ‘sunscreen’ due to their antioxidant capabilities, protecting the plant from excess sunlight derived damage (Gould et al. 2002; Araguirang and Richter 2022; Nowak et al. 2024). Nitrogen deficiency may trigger this response more readily, lowering the light intensity threshold at which anthocyanin biosynthesis is activated as a stress response. One reason for this role of nitrogen might be connected to photosynthesis. Low nitrogen levels lead to a downregulation of photosynthesis (Evans and Clarke 2019; Mu and Chen 2021). Due to the long stress application in the present experiment (35 days), the amount of available photosynthesis machineries in the plants under nitrogen deprivation under light conditions could have been reduced, resulting in a higher amount of excess light energy. While plants can be protected from an excess of light energy by mechanisms like non-photochemical quenching (NPQ), a downregulation of photosynthesis also diminishes such protective mechanisms (Müller et al. 2001; Ruban and Wilson 2021). Thus, the plants under nitrogen deprivation and light conditions might not have been able to cope with excess light without another protection: anthocyanins.

It remains open why particularly the traps seem to be the first part of *U. gibba* to turn red. Is it to attract prey, or is it to cope with an excess of supplied carbon (Sirová et al. 2010)? While the accumulation of anthocyanins has been reported to occur under a multitude of stress factors, it is shown here, that nitrogen deficiency alone does not elevate anthocyanin biosynthesis gene expression without the presence of light in *U. gibba*.

## Methods

### A. Plant material and growing conditions

*U. gibba* L. was provided by the Botanical Garden Darmstadt (XX-0-DATH-519) and surface sterilized by first, briefly rinsing with soapy dH_2_O, followed by an incubation with sterile-filtrated soapy 0.07% sodium hypochlorite for 60 seconds. Next, the plant material was briefly washed with sterile dH_2_O and placed on media. Stock cultures are grown in glass jars of 370 ml size (WECK-Sturzglas RR100) that were filled with ca. 75 ml 1/4 MS2 (one-quarter-strength Murashige and Skoog medium with 2% (w/v) sucrose) + 0.25 mg/l 6-benzylaminopurine (BAP) liquid media and divided and placed on fresh media every 2-3 weeks (Murashige and Skoog 1962). The conditions are ca. 2400-2700 lx 16/8 h day/night light (Philips TL5 39W/840 HO) and 22-25 °C. All plant cultures originate from *U. gibba* (XX-0-DATH-519) and were grown in *in vitro* cultures with ca. 2400-2700 lx (light) or ca. 5-10 lx (dark) 16/8 h day/night light (Philips TL5 39W/840 HO) and 22-25 °C in glass jars of 370 ml size (WECK-Sturzglas RR100) that were filled with ca. 75 ml Hoagland solution (HS), either normal or without nitrogen (HiMedia, REF TS1094-5L and TS1117-5L). From a stock culture grown in 1/4 MS2 + 0.25 mg/l 6-benzylaminopurine (BAP), cultures containing 1/4 MS2 without BAP were grown for 13 or 14 days. From these, 1-2 cm long pieces were cut and used to start cultures containing HS with or without nitrogen, placed in a way that the whole bottom of the new flask was slightly covered with plant material (ca. 20-30 pieces). These were placed in light or dark conditions. After 35 days, the cultures were harvested for RNA extraction. In total, per media condition (normal or without nitrogen), 5 samples with light and 4 samples in dark were used for RNA-seq (18 total).

### B. RNA extraction and RNA-seq

For RNA extraction, a culture was first washed with dH_2_O while kept in their culture vessel. Excess water was briefly removed by flipping the culture vessel while the lid was held canted. More thorough drying was carried out by pressing the plant material between two large layers of tissue paper before grinding it with a mortar and pestle with liquid nitrogen. Between pressing the plant material in tissues and coming in contact with liquid nitrogen, only up to 10 seconds passed. RNA extraction was carried out with Plant and Fungi RNA kit from Macherey-Nagel (REF 740120.50). Ca. 100-200 mg of plant material were used as sample input for step 1A and the kit instructions were followed from that on (03/2025, Rev. 07). The following exceptions were made from these instructions: Between first and second wash at step 5 a DNase digest was conducted on-column with 100 µl rDNase solution; final elution in 20 µl for most samples and 30 µl for some other samples (Macherey-Nagel, REF 740949.50) applied for 10 minutes at 37 °C. The remaining plant powder was stored at -70 °C without intermediate thawing. After measuring the RNA spectrophotometrically, it was stored at -70 °C. Paired-end RNA-seq was conducted by a third party using Illumina NovaSeq X Plus Series sequencing system (PE150).

### C. DNA extraction and sequencing

For genome sequencing, high molecular weight DNA was extracted from an *U. gibba in vitro* stock culture grown in BAP media following a previously developed CTAB protocol optimized for high molecular weight DNA extraction from plants (Siadjeu et al. 2020; Wolff et al. 2024; de Oliveira et al. 2026). Sequencing was performed on R10.4.1 flow cells. Library construction was performed using 1 µg of high molecular weight DNA following the SQK-LSK114 protocol. Priming of the flow cells and washing steps between sequencing runs were performed according to ONT’s protocols (EXP-FLP002, EXP-WSH004). Sequencing was performed on a MinION Mk1B. Base-calling of sequencing data stored in POD5 files was performed with dorado v0.8.3 (Oxford Nanopore Technologies, 2024).

### D. Genome sequence assembly and softmasking

Genome sequence assembly was generated with hifiasm (version: 0.25.0-r726), giving the raw reads as input and the ‘–ont’ flag (Cheng et al. 2024). From hifiasm’s results, the assembly graph of primary contigs (prefix.bp.p_ ctg.gfa) were converted into a FASTA file with awk ‘/^S/ {print “>“$2;print $3}’ (version: GNU Awk 5.2.1). Soft-masking was conducted by first, running EDTA in the provided docker container (quay.io/biocontainers/edta:2.2.2— hdfd78af_1) with the options ‘-anno 1 -sensitive 1’ and second, the EDTA script ‘make_masked.pl’ (from the 2.2.2 release) with the options ‘-minlen 80 -hardmask 0’ (Ou et al. 2019). This softmasked genome sequence is named daUtrGib.v01.

#### D.1. Genome sequence quality metrics

BUSCO score was calculated with the provided docker container (ezlabgva/busco:v6.0.0_cv1), using the lamiales_odb12 lineage dataset (Tegenfeldt et al. 2025a). LAI score was calculated using the EDTA results and the LAI from the same docker container version (see above) (Ou et al. 2018). Simple statistics (total size, N50, no. of sequences) were obtained with seqkit v2.11.0 (Shen et al. 2024).

### E. Structural annotation

Briefly, structural annotation was obtained by combining individual annotations from Helixer, BRAKER3, and GeMoMaPipeline, with GeMoMa Annotation Filter (GAF) (Holst et al. 2026; Gabriel et al. 2023; Keilwagen et al. 2018). The docker version of Helixer version 0.3.6 (gglyptodon/helixer-docker:latest) was applied to daUtrGib.v01 with the land_plant_v0.3_a_0080.h5 model and the ‘–subsequence-length 64152’ option. BRAKER3 version 3.0.8 was run with protein hints and RNA-seq hints, provided by a mapping (BAM file). Protein hints have been generated by combining the FASTA files of Viridiplantae odb12 with all UniProt entries that could be found for taxonomy_id 4143 (Lamiales) as of 2026-02-01 (see Additional file 10 for list of UniProt IDs) (The UniProt Consortium et al. 2025; Tegenfeldt et al. 2025b). RNA-seq reads of the INSDC projects PRJEB110519, PRJEB110496, and PRJEB111052 have been mapped with HISAT2 (version 2.2.1) to a HISAT2 index created from a non softmasked version of daUtrGibb.v01 and the STDOUT of the HISAT2 mapping command has been piped to “samtools view -h -F 4 -| samtools sort -m 2G - -O BAM -o .sorted.bam –write-index” to save the results into a sorted BAM file, create an index of it, and to save space by only saving mapped reads into it. The GeMoMa-1.9.jar was executed with the java parameters ‘-jar -Xmx200G’ to run the GeMoMa program ‘CLI GeMoMaPipeline with the parameters ‘pc=true pgr=true o=true Extractor.r=true GAF.f=“start==‘M’ and stop==‘*’ and (isNaN(score) or score/aa>=‘0.75’)” AnnotationFinalizer.r=NO’. Additional hints were the RNA-seq reads provided with the same BAM file as used for BRAKER3, specified with the options ‘r=MAPPED ERE.m=.sorted.bam’ and the genome sequences along with their structural annotation as GFF file from *U. gibba* (CoGe genome ID: 29027) (Lan et al. 2017), *U. reniformis* (Silva et al. 2019), *Pinguicula gigantea* (CoGe genome ID: 59509) (Fleck et al. 2025) and *Genlisea aurea* (NCBI ID: GCA_000441915.1) (Leushkin et al. 2013). Merging of the individual annotations was constructed by first running GeMoMa ERE on the RNA-seq reads BAM file (described above) to extract RNA-seq hints. Second, with GeMoMa AnnotationEvidence and the extracted RNA-seq hints, the annotations from BRAKER3 and Helixer have been annotated. Third, GeMoMa GAF was applied once to all of these three GeMoMa filterable annotations to obtain a final annotation. The filter options were f=“start==‘M’ and stop==‘*’ and aa>=15 and (isNaN(score) or score/ aa>=‘0.75’) and (evidence>1 or sumWeight>1 or avgCov>1 or tpc==1.0)”. To finalise the merged annotation, GeMoMa AnnotationFinalizer was run to the mergedGFF with the options ‘tf=true p=daUtrGibb_v01_v01’, to create a properly named annotation. To afterwards replace the underscores with dots, GNU sed 4.9 was applied. The GFF file was further cleaned for upload to ENA. First, AGAT (version: v1.6.1) agat_sp_fix_features_ locations_duplicated.pl and agat_convert_sp_gxf2gxf.pl were applied (Dainat 2022). Then, EMBLmyGFF3 (version: 2.4 pyhdfd78af_1 bioconda) was used with the parameters “–topology linear –molecule_type ‘genomic DNA’ –transl_table 1 –species ‘13748’ -i UGIBB -p PRJEB97978” to create a EMBL flat file which was gzip compressed with pigz (Norling et al. 2018). With the ENA Webin-CLI tool (version: 9.0.3) and the options ‘-validate -context=genome’, the files were first validated. A specific error was fixed with a custom script (clean_gff_of_ ENA-WEBIN-CLI_ERROR_Abutting_features_cannot_ be_adjacent_v03.py, https://codeberg.org/snmeckoni/scripts) and validation and fixing was conducted multiple times consecutively until a valid GFF file was obtained (daUtrGibb.v01.v01.gff) on which the uploaded genome sequence and annotation is based (GCA_982292365.1). On this GFF file, GeMoMa Extractor with the options ‘p=true c=true identical=true’ was applied to generate cds and pep FASTA files (daUtrGibb.v01.v01.pep.fasta, daUtrGibb.v01.v01.cds.fasta). AGAT agat_sp_keep_ longest_isoform.pl was applied to daUtrGibb.v01.v01.gff to generate daUtrGibb.v01.v01.longest.isoforms.gff and subsequently GeMoMa Extractor to obtain daUtrGibb.v01.v01.longest.isoforms.pep.fasta and daUtrGibb.v01.v01.longest.isoforms.cds.fasta.

### F. Functional annotation

For daUtrGibb.v01.v01.longest.isoforms.pep.fasta, proteins associated with the anthocyanin metabolism were identified with two dedicated tools. To identify the structural genes of the flavonoid biosynthesis pathway, an analysis with KIPEs v3.2.6 (Rempel et al. 2023) and the flavonoid baits data set v.3.4 was conducted. Flavonoid biosynthesis controlling MYB transcription factors were annotated using the MYB_annotator v1.0.3 (Pucker 2022) with parameters described in Additional file 11. For daUtrGibb.v01.v01.pep.fasta, a general annotation was produced using construct_anno.py and the functional annotation available for *A. thaliana* (Pucker and Iorizzo 2023; Cheng et al. 2017).

### G. RNA-seq analyses

The raw RNA-seq reads have been processed with kallisto (version 0.52.0) against all and only the longest isoforms of the predicted genes from *U. gibba* (Bray et al. 2016). First, a kallisto index was created with the kallisto index command and the respective CDS FASTA file. Second, kallisto quant was executed for each sample individually with the script iterate_kallisto_quant_03.sh and the results were merged with the script kallisto_subfolder_merger_v02.sh (both scripts are available from a codeberg.org repository, release v.1.1, archived at Zenodo). DESeq2 was applied to the kallisto results with an R script to generate statistics per gene and a PCA plot (Love et al. 2014). For each lighting condition the relation between the two different media conditions was calculated, as well as the interaction between those. The script, input files, and output files can be obtained from the corresponding codeberg.org repository under the folder DESeq2_analysis (repository release v1.1, archived at Zenodo. From the KIPEs results, best hits have been manually curated and used to create an overview heatmap (Additional file 6, corresponding additional data (script and input files) is available in the folder additional_ file_6 from the corresponding codeberg.org repository, release v.1.1, archived at Zenodo). From that, relevant genes have been chosen for a refined heatmap focusing on the relevant anthocyanin biosynthesis genes (Figure 2), including the DESeq2 results as bar plots (additional data (script and input files) is available in the folder figure_2 from the corresponding codeberg.org repository, release v.1.1, archived at Zenodo). Figure 2 was additionally edited with Inkscape. For the MYB_annotator results, genes of interest have been extracted with grep -e “phenylpropanoid” -e “anthocyanin” -e “flav” Additional_file_7_03b_new_2_ref_myb_ mapping_file.txt > filtered.MYB_candidates.list.txt. Combined with the DESeq2 and kallisto results, a heatmap has been created and manually edited and saved as JPEG with Inkscape (Additional file 8, additional data (script and input files) is available from the corresponding codeberg.org repository, release v.1.1, archived at Zenodo).

## ACKNOWLEDGEMENTS

We thank the Plant Biotechnology and Bioinformatics group members and the Botanic Gardens Bonn staff. We thank Simon Poppinga for providing initial plant material, America Cox and Tim Katzek for the help with the experiments, and Samud Suhas Shetty for proof-reading the manuscript. Claude LLM was used to enhance the R script with the option to use positional arguments and to add the barcharts plotting function to the heatmap python scripts.

## Supplementary Information

**Additional file 1:** Photograph of an *U. gibba* control sample under light conditions. The picture was taken right before harvest and is presented in JPEG file format.

**Additional file 2:** Photograph of an *U. gibba* control sample under dark conditions. The picture was taken right before harvest and is presented in JPEG file format.

**Additional file 3:** Photograph of an *U. gibba* nitrogen deprivation sample under light conditions. The picture was taken right before harvest and is presented in JPEG file format.

**Additional file 4:** Photograph of an *U. gibba* nitrogen deprivation sample under dark conditions. The picture was taken right before harvest and is presented in JPEG file format.

**Additional file 5:** Result table of anthocyanin biosynthesis genes, functionally annotated in *U. gibba* with KIPEs based on daUtrGibb.v01.v01.longest.isoforms.pep.fasta. The file is presented in TXT format.

**Additional file 6:** Expression pattern of candidate structural flavonoid biosynthesis genes under nitrogen starvation in the presence or absence of light. The heatmap is based on kallisto TPMs, and the color is z-score normalized by gene/row. Rows are labeled by the daUtrGibb.v01.v01 gene ID and its corresponding KIPEs annotation, as well as the conserved residue score assigned by KIPEs. The file is presented in PNG format.

**Additional file 7:** Result table of MYB transcription factor genes, functionally annotated in *U. gibba* with KIPEs based on daUtrGibb.v01.v01.longest.isoforms.pep.fasta. The file is presented in TXT format.

**Additional file 8:** Expression pattern of candidate MYB transcription factor genes under nitrogen starvation in the presence or absence of light. The heatmap is based on kallisto TPMs, and the color is z-score normalized by gene/ row. The bar charts next to the heatmap show the log2 fold change between nitrogen containing and nitrogen starvation media for normal light conditions (orange bars, L) and dark conditions (blue bars, D). If the change in gene expression between the different media conditions is significant (padj ≤ 0.05), the bar is enriched by an arrow. The bracket with an asterisk marks a significant difference (padj ≤ 0.05) in the log2 fold change between the different light conditions (DESeq2 interaction). Rows are labeled by the daUtrGibb.v01.v01 gene ID and its corresponding MYB_annotator annotation, as well as their putative function assigned by the MYB_annotator. Row labels have been colored according to the corresponding genes belonging into subgroup 6, 7, or 4 MYBs based on the annotation. The file is presented in JPEG format.

**Additional file 9:** Table of daUtrGibb.v01.v01 gene IDs and their corresponding structural anthocyanin biosynthesis genes, manually curated based on the KIPEs results.

**Additional file 10:** List of UniProt IDs of Lamiales (taxonomy_id 4143) reference proteins used for structural gene annotation (as of 2026-02-01) as TXT format file.

**Additional file 11:** Documentation of the MYB_annotator analysis as TXT format file.

## Data Availability

Raw reads are available in INSDC via project PRJEB97978 (genomic reads, genome assembly, and annotation) and PRJEB110519, PRJEB110496, and PRJEB111052 (RNA-seq data). Scripts and detailed input files are available via a codeberg.org repository (release v.1.1 (https://codeberg.org/snmeckoni/daUtrGibb.v01_additional_data_repository/releases/tag/v.1.1), archived at Zenodo (https://doi.org/10.5281/zenodo.22119014). Assembly and annotation files for daUtrGibb.v01.v01 are available at bonndata: https://doi.org/10.60507/FK2/JJ5QZX.

