## Supplementary figures and images for "Anthocyanin biosynthesis gene activation in nitrogen deprived *Utricularia gibba* L. under light or darkness"

### Additional file 1

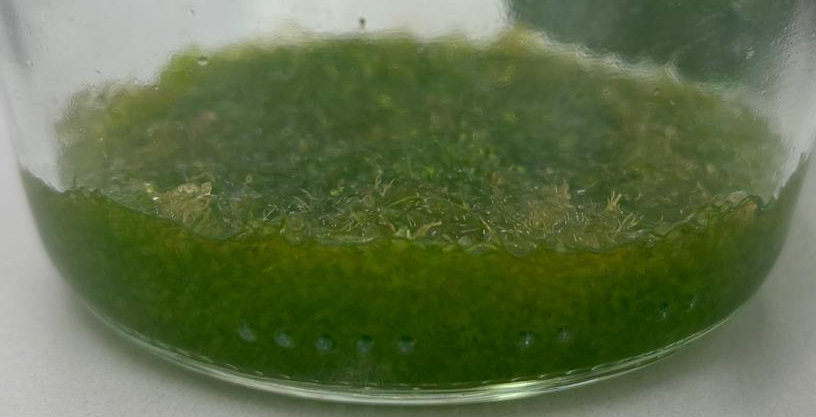

### Additional file 2

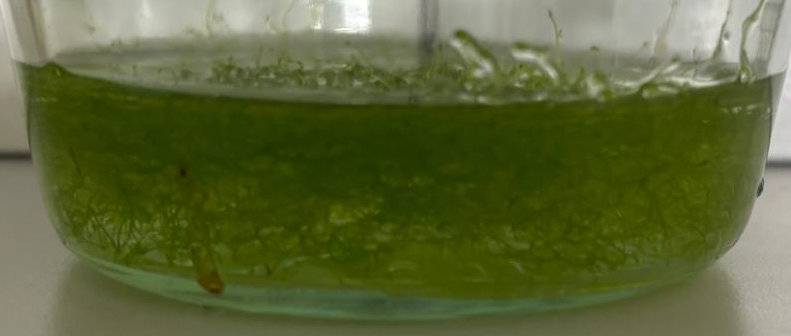

### Additional file 3

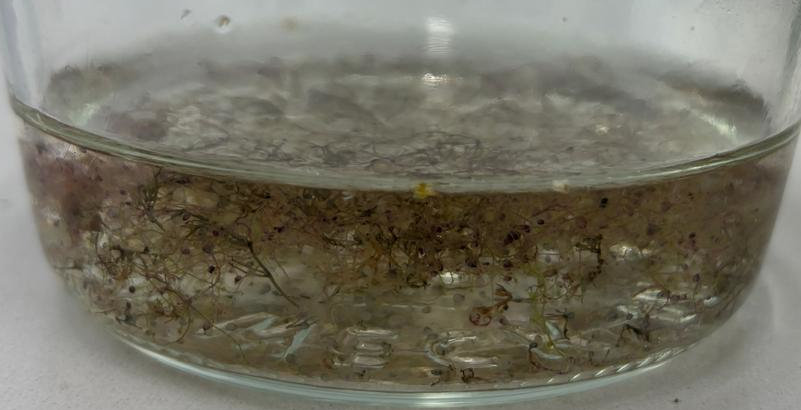

### Additional file 4

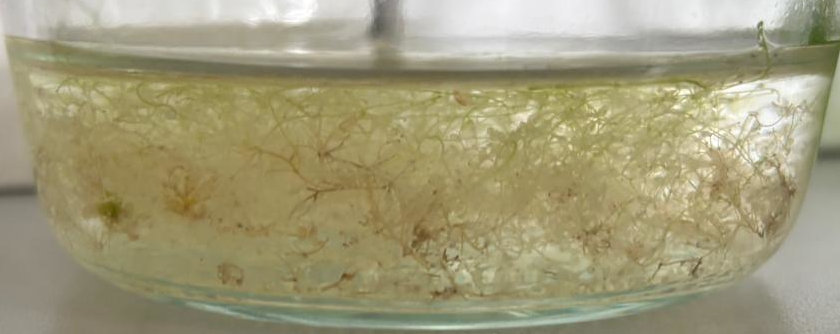

### Additional file 6

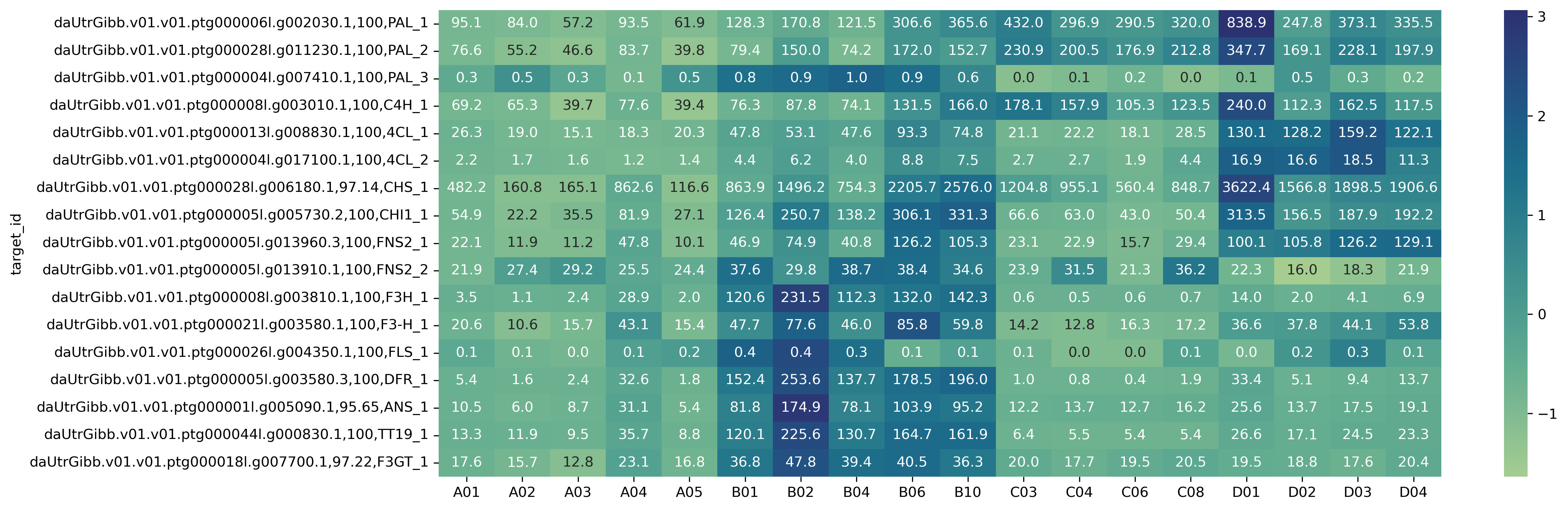

### Additional file 8

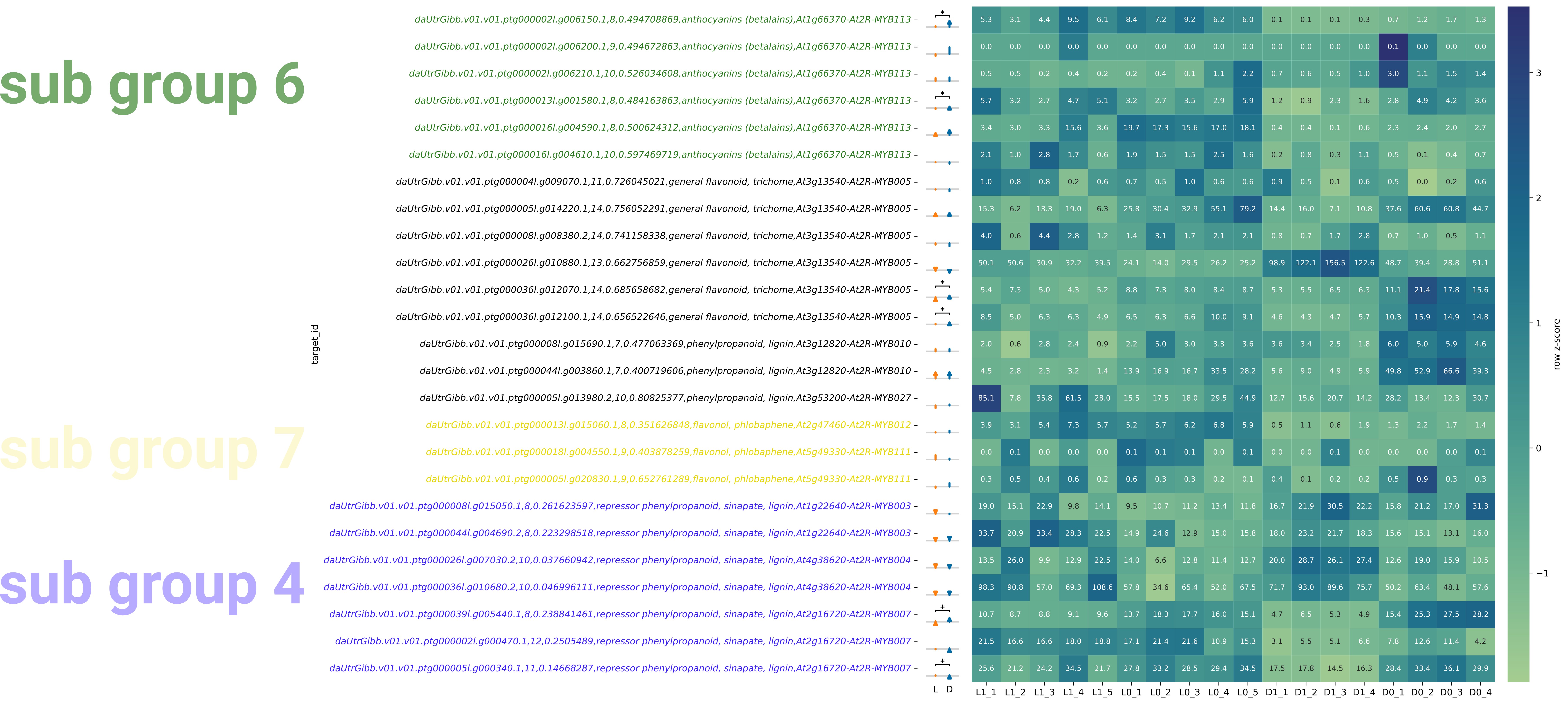
