## Additional file 11 for "Anthocyanin biosynthesis gene activation in nitrogen deprived *Utricularia gibba* L. under light or darkness"

Please cite 'Pucker B (2022). Automatic identification and annotation of MYB gene family members in plants. BMC Genomics 23, 220 (2022). doi: 10.1186/s12864-022-08452-5' when using MYB_annotator.py.

MYB_annotator.py version: v1.0.3
MYB bait file: /vol/data/tools/MYB_annotator/MYB_baits.fasta    f92f35b0d351312c43a6e1b30aa0567e
MYB info file: /vol/data/tools/MYB_annotator/MYB_baits.txt      5836d7f542b672365883fa0a29d5d5a0
Subject FASTA file: /vol/data/daUtrGibb.v01/daUtrGibb.v01.v01.longest.isoforms.pep.fasta        2beb0d582587a49762d68e5d755e814e
Output folder: /vol/data/XXX
Tool for initial candidate selection: blast
Tool for tree construction: fasttree
CPUs for BLASTp: 13
CPUs for RAxML: 13
Reference MYB file: /vol/data/tools/MYB_annotator/AthRefMYBs.txt        abc8047bebc188b8a20935fe82467b5d
Type of input: PEP
blastp path: blastp
makeblastdb path: makeblastdb
mafft path: mafft
raxml path: raxml-ng
fasttree path: fasttree
Minimal BLASTp hit similarity cutoff: 60.0
Maximal number of BLASTp hits per bait: 100
Minimal BLASTp hit alignment length: 75
Number of neighbours to consider in classification: 10
Factor for branch length cutoff in identification of neighbours: 10.0
Minimal number of neighbours required for classification as MYB: 0
Distance cutoff for paralog clade masking: 10.0
MAFFT version: v7.525 (2024/Mar/13)
FastTree version:  2.2.0 D
RAxML version: RAxML-NG v. 2.0.0 released on 11.03.2026 by The Exelixis Lab....
hmmsearch version detection failed.
